# Denisovan introgression has shaped the immune system of present-day Papuans

**DOI:** 10.1101/2020.07.09.196444

**Authors:** Davide M. Vespasiani, Guy S. Jacobs, Laura E. Cook, Nicolas Brucato, Matthew Leavesley, Christopher Kinipi, Francois-Xavier Ricaut, Murray P. Cox, Irene Gallego Romero

## Abstract

Modern humans have substantially admixed with multiple archaic hominins. Papuans, in particular, owe up to 5% of their genome to Denisovans, a sister group to Neanderthals, whose remains have only been identified in Siberia and Tibet. Unfortunately, the biological and evolutionary significance of these introgression events remain poorly understood. Here we investigate the function of archaic alleles of both Denisovan and Neanderthal ancestry characterised within a previously published set of 56 genomes from individuals of Papuan genetic ancestry living in the island of New Guinea. By comparing the distribution of archaic and modern human variants, we are able to assess the consequences of archaic admixture across a multitude of different cell types and functional elements. We detect a consistent signal across Denisovan variants of strong involvement in immune-related processes throughout our analyses. Archaic alleles are often located within cis-regulatory elements and transcribed regions of the genome, suggesting that they are capable of contributing to a wide range of cellular regulatory processes. We identify 3,538 high-confidence Denisovan variants that fall within annotated cis-regulatory elements and have the potential to alter the affinity of multiple transcription factors to their cognate DNA motifs, highlighting a likely mechanism by which introgressed DNA can impact phenotypes in present-day humans. Lastly, we experimentally validate these predictions by testing the regulatory potential of five Denisovan variants segregating at high frequency within Papuan individuals, and find that two are associated with a significant reduction of transcriptional activities in plasmid reporter assays relative to modern human alleles. Together, these data provide support for the hypothesis that, despite their broadly deleterious nature, archaic alleles actively contribute to modern human phenotypic diversity to this day, and might have facilitated early adaptation to non-African environments.

## Introduction

Modern humans are known to have interbred with Neanderthals [1], Denisovans [2] and possibly other archaic hominins [3]. While genetically similar populations of Neanderthals are thought to have contributed approximately 2% to non-African genomes, Denisovan introgression has been observed to be more variable [4]. Particularly, Denisovan ancestry accounts for up to 5% of the genomes of the Indigenous peoples of Island Southeast Asia and Australia [4, 5]. In addition, these components exhibit a deep divergence from the reference Altai Denisovan genome, providing strong evidence for the occurrence of multiple Denisovan introgression events across time and space [6, 7].

At the genomic level these introgressed archaic alleles are mostly observed outside protein-coding sequences, distributed over non-functional and regulatory regions [8, 9]. Enhancers, in particular, are amongst the top targeted elements of archaic hominin introgression [10,11]. Here, archaic alleles are thought to drive phenotypic differences by altering gene pre- and post-transcriptional regulatory processes [11]. Furthermore, both Neanderthal and Denisovan variants seem to preferentially affect enhancers in a tissuespecific manner, with highly pleiotropic elements being depleted of archaic variation [12]. However, beyond general agreement that introgressed archaic DNA has mainly been deleterious and actively removed from coding sequences and conserved non-coding elements [8, 12–14], the actual phenotypic consequences of this variation are not well understood. Several lines of evidence highlight associations between archaic DNA and risk for disease traits, including autoimmune diseases [8, 15, 16], or with traits of possible evolutionary advantage for early non-Africans [17, 18]. For example, Neanderthal variants within immune genes and immune-related cis-regulatory elements (CREs) have been associated with differential responses to viral infections among present-day Europeans [19, 20].

Unfortunately, two main factors limit our interpretation of the biology of these alleles. First, we still lack a detailed characterisation of global levels of human genetic diversity, vital to identifying differential archaic hominin contributions across modern populations [21, 22]. Because Denisovan DNA is not present in the genomes of modern Europeans, which make up the vast diversity of biomedical genetics cohorts, it has been impossible to link them to phenotype, unlike segregating Neanderthal variants [16, 23]. Second, and more challenging, most of these alleles lie within non-coding sequences where, despite their acknowledged contributions to human evolutionary history [24, 25], an understanding of their actual biological functions remains elusive.

To gain insights into the consequences of archaic introgression in Papuans, we have analysed a previously published dataset of 56 present-day genomes sampled across the island of New Guinea [6]. By comparing the distribution of archaic single nucleotide polymorphisms (aSNPs) and non-archaic SNPs (naSNPs) segregating within these populations, across multiple genomic elements and cell types, we find that aSNPs are enriched within functional cis-regulatory and transcribed elements, particularly those active within immune-related cells. We also observe that the presence of archaic alleles within these elements can lead to substantial disruption of the binding sites of transcription factors, consistently pointing towards an involvement of Denisovan variants in immune processes. We validate this through experimental testing of multiple Denisovan aSNPs, and find they are associated with significant transcriptional changes in reporter gene experiments.

## Results

### Building a high-confidence set of archaic variants and SNPs annotation

To curate list of high-confidence archaic SNPs to analyse for possible functional contributions to present-day Papuans, we took advantage of a recently described dataset of both archaic and non-archaic haplotypes segregating within 56 individuals of Papuan ancestry [6]. Due to incomplete lineage sorting and recombination, a large fraction of variants included in these archaic haplotypes is not expected to be of archaic origin [9]. To account for this while enriching for variants of evolutionary interest we applied a series of filtering steps to our starting list of 7,791,042 genomic SNPs from [6], classified as either archaic (aSNPs, segregating within either Denisovan or Neanderthal introgressed haplotypes) or non-archaic (naSNPs, segregating in all other haplotypes). A series of stringent filtering steps (see Methods) resulted in three high confidence sets of 79,808, 51,268 and 338,585 variants which we assigned to respectively to Denisovan, Neanderthal and modern (postdating the Out of Africa event) human ancestries.

### Distribution of aSNPs across chromatin states and tissues

Given that both aSNPs and naSNPs mainly lie outside coding regions [8, 9] we decided to compare the distribution of aSNPs and naSNPs across chromatin functional states and human cell types. Thus, we examined the distribution of our variants against epigenomic maps of 15 different chromatin states across 111 cell types generated by the Roadmap Epigenomics Project [26]. To identify chromatin states harbouring a significant excess or deficit of archaic variants, we counted the number of aSNPs and naSNPs that fall within a given chromatin state in at least one cell type (i.e., without controlling for pleiotropy) and computed the odds ratio (OR) between aSNPs and naSNPs. We found that aSNPs are generally depleted from transcription start sites (TssA) and from the majority of inactive chromatin states (i.e. states not associated with gene expression, as defined by [26]). The exception to this trend was quiescent chromatin, which consistently harboured the largest number of variants. This is expected given this state is predominant in each epigenome [26], and suggests that a substantial fraction of aSNPs might be functionally inert (Fig. 1).

**Fig 1.**
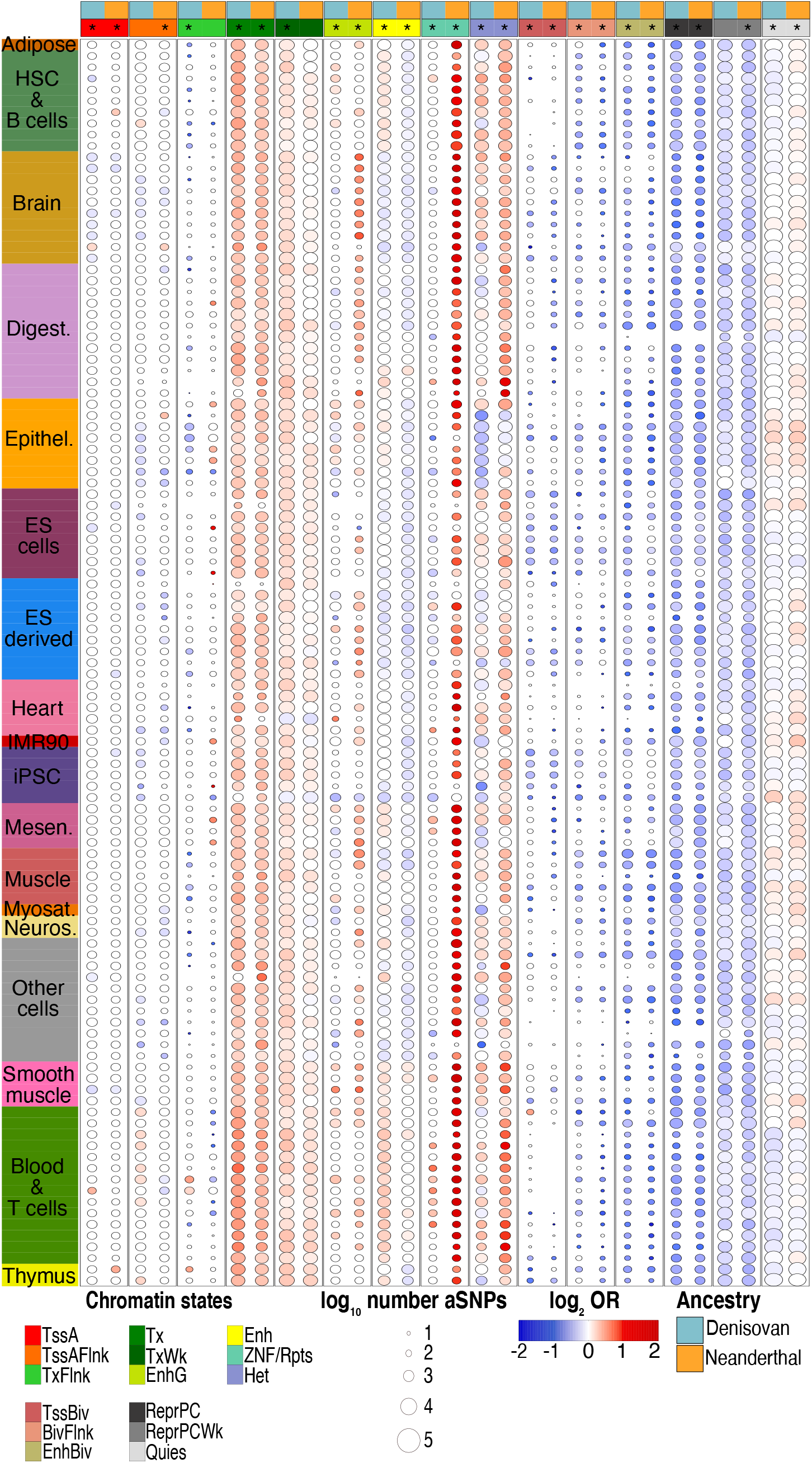
aSNP introgression across the Roadmap Epigenomics data. Heatmap of the Denisovan and Neanderthal aSNPs impact across each chromatin state/cell type combination. Only significant differences (i.e. FDR-corrected Fisher’s exact test *p* < 0.05 are coloured). Asterisks indicate genome-wide significant enrichment/depletion of aSNPs for the given state (see Fig. S1 and Supp Table 1).

Both Denisovan and Neanderthal aSNPs were significantly enriched within transcribed states (Tx), genic enhancers (EnhG) and, especially in the case of Neanderthals, within zinc-finger protein-genes and repeats (ZNF/Rpts) states. In addition, Denisovan, but not Neanderthal aSNPs, were significantly enriched within weakly transcribed regions (TxWk), enhancers (Enh) and significantly depleted from heterochromatin (Het) and from TxFlnk, a state showing both enhancer- and promoter-like epigenetic marks [26]. Conversely, Neanderthal variants were significantly enriched within Het, but significantly depleted from Enh and promoters (TssAFlnk) (Fig. S1 and Supp Table 1).

We then sought to understand whether these observations were consistent across the entire Roadmap Epigenomics dataset, or driven by a subset of cell types. Denisovan aSNPs were significantly depleted from quiescent regions within haematopoetic stem cells (HSC) & B cells, blood & immune T cells and thymus, whereas Neanderthal depletion was less consistent (Fig. 1 and Supp Table 1). Likewise, Denisovan aSNPs were strongly over-represented within Enh and EnhG in HSC & B cells, blood & immune T cells and thymus, tissues and states from which Neanderthal variants were largely depleted. On the other hand, only Neanderthal aSNPs were strongly enriched within EnhG states active in brain, digestive tissues as well as within a large number of toti/pluripotent cell types (Fig. 1 and Supp Table 1). The strongest enrichment signal for Neanderthal aSNPs was within ZNF/Rpts, which were in excess across almost all cell types. This state spans regions of the genome containing actively transcribed ZNF genes, and is characterised both by H3K36me3 (active transcription) and H3K9me3 (heterochromatin associated) epigenetic marks as well as highly enriched for satellite repeats [26], making evaluation of this finding challenging.

Finally, to confirm whether these signals were caused by SNPs occurring within highly pleiotropic or cell-specific functional elements, we counted the number of different cell types across which a/naSNP-containing elements were annotated with any given chromatin state. This yielded an estimate of the potential pleiotropic activity of each functional element. We detected similar degrees of pleiotropy across all the three ancestries, with variants annotated within Quies, TssA, Tx and TxWk exhibiting higher pleiotropic activities than those annotated within TssAFlnk, TxFlnk, Enh and EnhG, in line with findings by [26] (Fig. S2). Taken together, these findings suggest that aSNPs are likely to occur and be maintained within constitutive states that are functionally inert (Quies), perhaps as a consequence of purifying selection being reduced in those regions [27]. At the same time, our results indicate that a non-trivial fraction of introgressed variants still segregating within Papuans today might have functional consequences, particularly for gene regulatory processes, in a limited number of cell types.

Given our observations of the potential tissue specificity of archaic regulatory variants, we sought to investigate the effects of Denisovan and Neanderthal aSNPs on gene regulation. Hence, we restricted our analyses to all SNPs annotated within states associated with cis-regulatory elements (CREs, i.e. TssA, TssaFlnk, TxFlnk, Enh and EnhG). To prioritise variants of potential evolutionary importance we only retained SNPs with allele frequency > 0.2 (see Methods), as these might be under positive selection in our target population. This resulted in three sets of 8,548, 7,198 and 58,828 Denisovan, Neanderthal and modern human high-frequency variants respectively (46.7%, 44.9% and 47.6% of the set of all variants with allele frequency > 0.2, respectively).

We first intersected these variants with significant eQTLs (qval < 0.05) from GTEx v8 [28]. Only 24 Denisovan, 50 Neanderthal and 515 modern human variants (0.28%, 0.69% and 0.87% of testable SNPs, respectively) were eQTLs in GTEx. To ensure these results were not biased towards variants that might be at high frequency only in Papuans, we repeated the analysis using all SNPs in our dataset. This time we found 82 Denisovan, 169 Neanderthal and 1,248 modern human SNPs (0.10%, 0.33% and 0.37% of the entire set of SNPs, respectively) within GTEx. Given the well-recognised bias towards individuals of European ancestry within the GTEx dataset, these findings both support the need to include under-represented populations into genetic surveys, and reaffirm that the majority of our introgression assignments are correct, and, as expected, that our variants are geographically restricted. Moreover, the higher fraction of cis-eQTLs found in the subset of CRE-associated variants suggests our filtering strategy is effective at enriching for likely regulatory variants.

### Denisovan aSNPs might alter gene regulation in immune-related cells

Given the observed enrichment within CRE-associated chromatin states, particularly for Denisovan variants, we next sought to characterise the potential for our variants (both modern and archaic) to alter gene regulation, and functional mechanisms by which this may occur. To this end, we examined the potential of high-frequency variants within CREs to disrupt known transcription factor binding sites (TFBSs), assessing their impact on 690 different position weight matrices (PWMs) drawn from the from HOCOMOCO v.11 [29] and Jaspar2018 [30] databases. After consolidating variants predicted to disrupt the same motif across the two databases (see Methods), we found that 3,538 Denisovan, 2,886 Neanderthal and 24,747 modern human variants (41.4%, 40.1% and 42.1% of testable variants, respectively) were predicted to disrupt at least one DNA binding site (Supp File 1-Supp File 3).

Next we assessed whether this subset of TFBS-disrupting aSNPs and naSNPs were equally represented across tissues. Both Denisovan and Neanderthal variants were significantly depleted from iPSCs and ESCs; Denisovan variants were also depleted from ESC-derived cells and brain. However, only Denisovan TFBS-disrupting aSNPs were over-represented across a suite of tissues: adipose (OR = 1.14, Fisher’s exact test *p* = 0.009), HSC & B cells (OR = 1.11, Fisher’s exact test *p* = 0.0035), blood & immune T cells (OR = 1.27, Fisher’s exact test *p* = 2.69 × 10^-9^) and thymus (OR = 1.14, Fisher’s exact test *p* = 0.009). (Fig. 2 and Supp Table 1).

**Fig 2.**
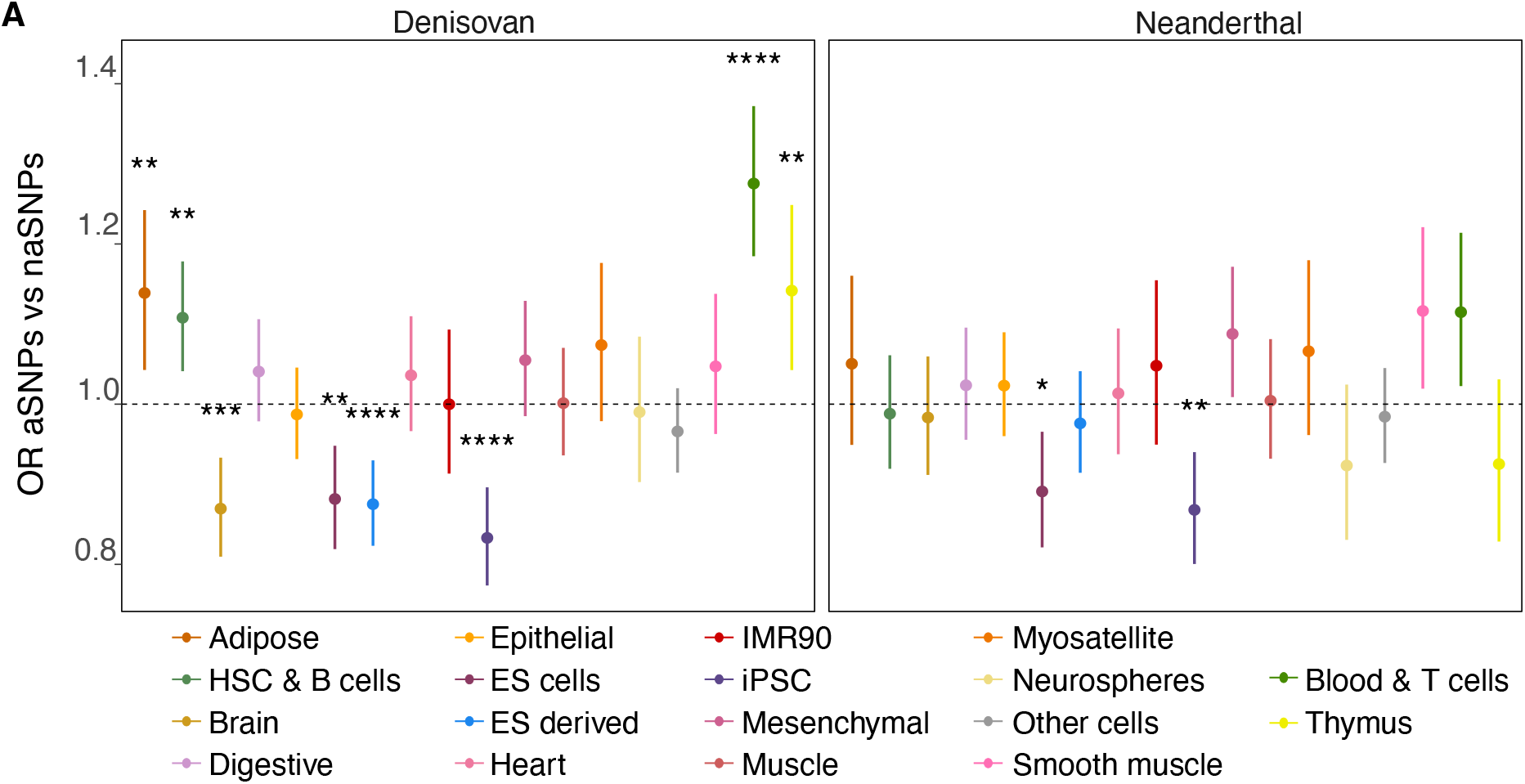
Impact of high-frequency TFBS-disrupting aSNPs across tissues and CRE-associated chromatin states. Denisovan and Neanderthal aSNPs enrichment across the 18 tissues and CRE-associated states. Dots and cross bars respectively represent the OR and upper and lower values. Asterisks indicate FDR-corrected Fisher’s exact test *p* < 0.05 (see Supp Table 1).

We next focused on the specific TFs driving these observations. To avoid redundancy due to similarities between predicted motifs across closely related TFs, we took advantage of recent work by Vierstra *et al*. [31], which clusters HOCOCOMO and Jaspar PWMs into 286 distinct clusters on the basis of sequence similarity, and considered only those a/naSNPs that disrupted PWMs included in these clusters (n = 3,028 Denisovan, 2,457 Neanderthal and 21,104 modern human variants). There were no genome-wide significant differences in the set of disrupted clusters between archaic and modern human variants (Fig. S3). Additionally, we sought to quantify the degree of disruption of the TFBS associated with each variant. To this end, we calculated Δ PWM, the difference in the PWM score between modern or introgressed alleles. All three ancestries contain SNPs predicted to have a significant disrupting impact on DNA motifs across multiple tissues, with no significant differences in the distribution of Δ PWM scores between aSNPs and naSNPs across tissues (Supp. Fig. S4 and Supp Table 2). Altogether, these findings suggest that archaic alleles do not preferentially disrupt specific DNA motifs nor tend to have significantly different impact compared to modern human variants. However, certain tissues might still harbour an excess of aSNPs that could also be associated with an unique set of disrupted TF clusters, potentially affecting tissue-specific gene regulatory processes.

### Denisovan TFBS-disrupting SNPs active in immune cells are mostly Papuan-specific

Recent studies have shown the existence of genetic structure characterising the Indonesian archipelago, with Papuan—and thus, Denisovan—ancestry showing a marked west to east cline [6, 32, 33]. We therefore investigated whether our entire subset of TFBS-disrupting variants also showed different frequencies between different human groups. To this extent, we defined aSNPs and naSNPs in genotype data from 63 individuals from western Indonesia, also part of [6], as above, and compared allele frequencies between Papuans and western Indonesians. 2,387 and 18,493 variants respectively of Neanderthal and modern human ancestry were shared between the two populations (82.7% and 74.7%, respectively), but this number falls to 1,683 (47.6%) for Denisovan aSNPs (Fig. 3a). Moreover, we found that shared Denisovan TFBS-disrupting variants segregate at very low frequencies in west Indonesians, whereas Neanderthal variants have higher variability (Fig. 3b).

**Fig 3.**
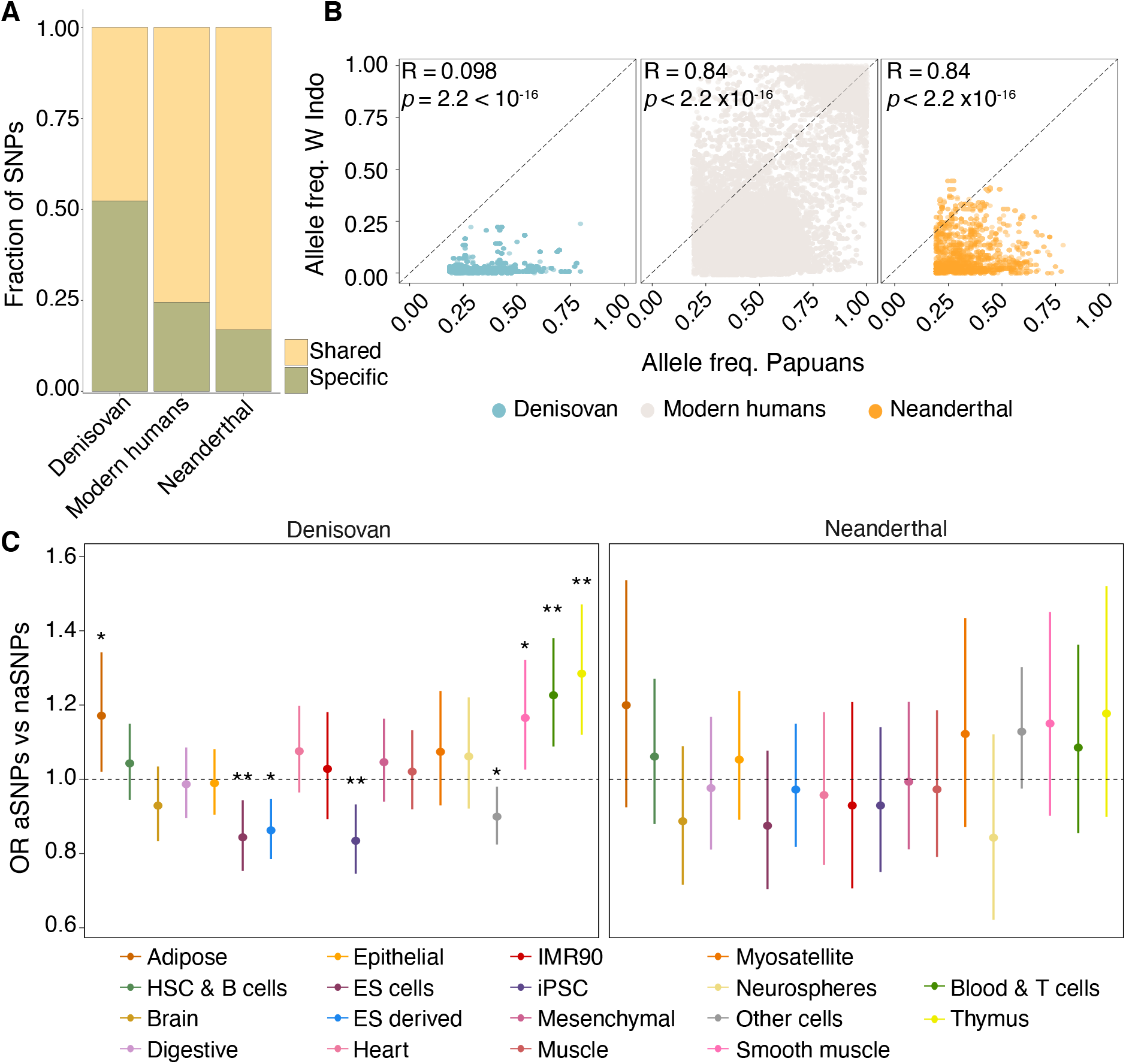
Distribution of TFBS-disrupting variants between western Indonesian and New Guinean populations. A) Proportion of variants shared across the Indonesian archipelago. B) Distribution of allele frequencies between western Indonesia and PNG for the subset of shared variants. C)Enrichment of Papuan-specific aSNPs across tissues. Asterisks indicate FDR-corrected Fisher’s exact test *p* < 0.05 (see Supp Table 1)

For each ancestry we then separately counted the Papuan-specific TFBS-disrupting SNPs annotated in each tissue and computed the OR between the number of Denisovan/Neanderthal and modern human variants. As expected, the subset of Neanderthal TFBS-disrupting SNPs active across any tissue was not significantly enriched for Papuan-specific variants relative to the set of modern human naSNPs. Conversely, we found a significant enrichment for Papuan-specific Denisovan TFBS-disrupting aSNPs within adipose (OR = 1.18, Fisher’s exact test *p* = 0.034), smooth muscle (OR = 1.16, Fisher’s exact test *p* = 0.034) thymus (OR = 1.29, Fisher’s exact test *p* = 0.0011) and blood & T cells (OR = 1.24, Fisher’s exact test *p* = 0.0011) (Fig. 3c).

### Denisovan TFBS-disrupting variants are predicted to affect immune-related processes

Given the predicted large impact of aSNPs, especially Denisovan, on immune-related cells, we next focused on the subset of high-frequency TFBS-disrupting SNPs annotated within CRE-associated states active in HSC & B cells and blood & immune T cells, retaining 1,481, 1,076 and 8,958 Denisovan, Neanderthal and modern human variants (41.8%, 37.3% and 36.2% of high-frequency TFBS-disrupting SNPs respectively). We then used GREAT [34] to link these variants to likely target genes, finding limited overlap in the sets of putative target genes between Denisovan and Neanderthal aSNPs (15 genes) or across all three ancestries (21 genes) (Fig. 4a). As expected, both aSNPs and naSNPs were distantly located from the nearest gene, with no differences among ancestries, further suggesting they lie within potential enhancer-like elements (Fig. S5).

**Fig 4.**
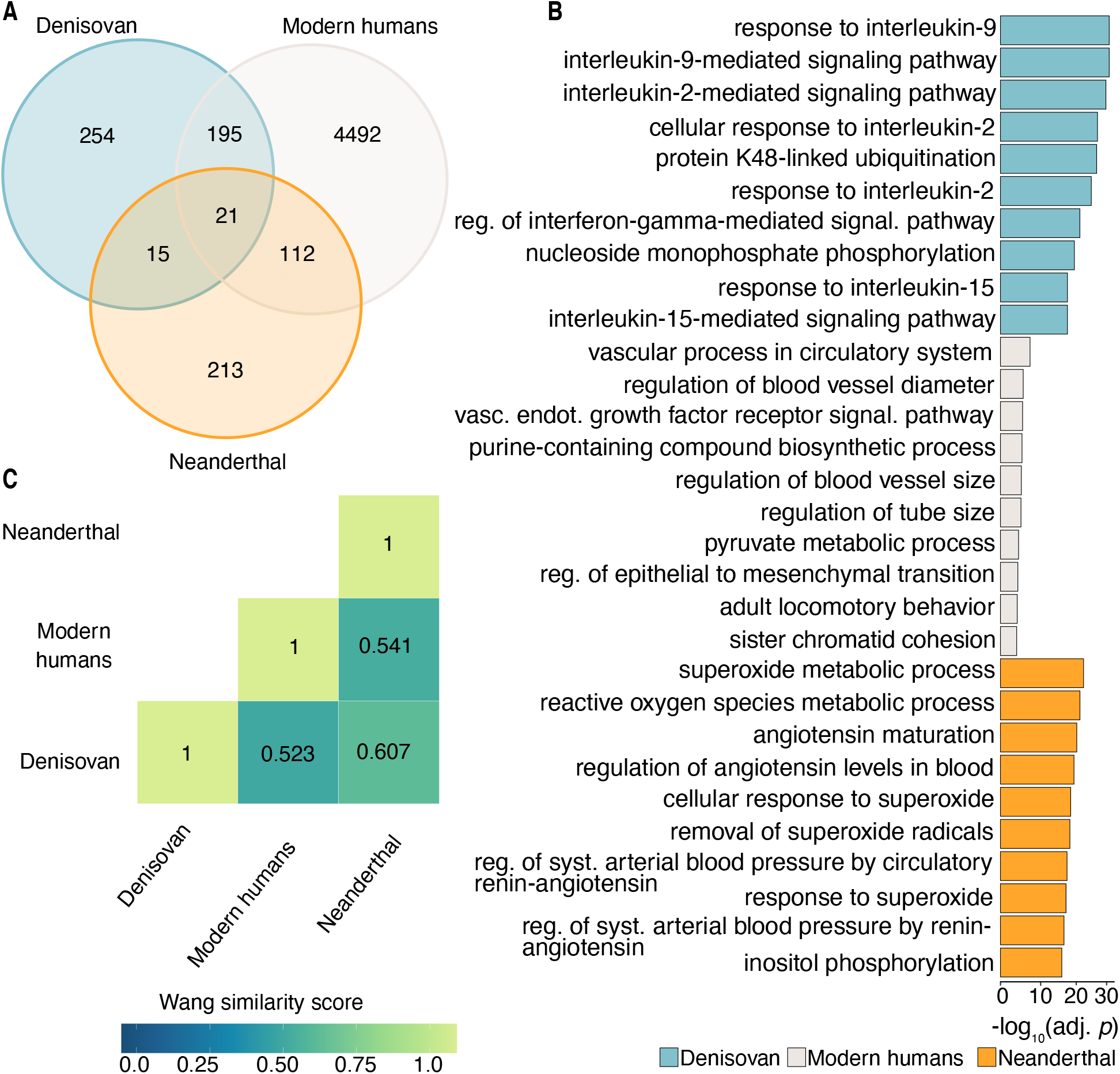
Biological processes possibly affected by TFBS-disrupting variants active in immune-related cells. A) Venn diagram with the overlap between the putative sets of target genes for each ancestry. B) The top 10^th^ mostly enriched GO terms for each ancestry. C) The semantic similarity score estimates between significantly enriched GO terms for each comparison.

We next performed Gene Ontology (GO) enrichment analyses on the set of genes associated with each ancestry. We found that Denisovan aSNPs, but neither Neanderthal aSNPs nor naSNPs were associated with genes strongly involved in multiple immune-related processes. Genes targeted by modern human and/or Neanderthal variants were instead enriched for more general biological processes, albeit instances related to neutrophil/granulocyte migration and chemotaxis were observed for Neanderthal variants (Fig. 4b; a full list of results is available as Supp Table 3). We then computed the semantic similarity between significantly enriched terms (FDR-adjusted hypergeometric *p* ≤ 0.01) using a method that incorporates both the locations of the terms in the GO graph as well as their relations with their ancestor terms [35], and found lower similarity between archaic and modern human enriched GO terms, while the similarity between Denisovan and Neanderthal enriched GO terms was the highest (Fig. 4c). Finally, for each ancestry we manually examined the set of genes associated with the significant GO terms. Denisovan TFBS-disrupting variants are predicted to regulate genes such as *TNFAIP3, OAS2* and *OAS3*, all of which have been repeatedly identified as harbouring archaic hominin contribution that impact immune responses to pathogens [36–38]. In particular, we found eight Denisovan TFBS-disrupting aSNPs (i.e., rs368816473, rs372433785, rs139804868, rs146859513, rs143462183, rs370655920, rs375463218 and rs372139279) associated with *OAS2* and *OAS3*. Notably, in all cases archaic alleles segregated at frequencies between 0.2 and 0.4 in Papuans, whereas west Indonesians were fixed for the reference, non-archaic alleles, and a comparison with 6 different non-human primate genomes indicated that for 7 of these 8 Denisovan aSNPs the introgressed allele is derived.

### Denisovan alleles are associated with regulatory changes in immune cells

The *OAS* locus has been repeatedly found to harbour signals of both Neanderthal and Denisovan adaptive introgression, with archaic alleles predicted to alter gene regulation [37, 38]. Our previous work has shown that both *OAS2* and *OAS3* are differentially expressed in whole blood between the people of Mentawai, a small barrier island off the coast of Sumatra, in West Indonesia, and the Korowai, a genetically Papuan hunter-gatherer group living on the Indonesian side of New Guinea island (Fig. 5b) [39]. To understand whether Denisovan introgression might contribute to these differences we tested the regulatory activity of 5 of the Denisovan variants described above (rs372433785, rs139804868, rs146859513, rs143462183 and rs370655920), all of which were predicted to disrupt the binding sites of TFs active in human immune cells, using a reported plasmid construct (see Methods).

**Fig 5.**
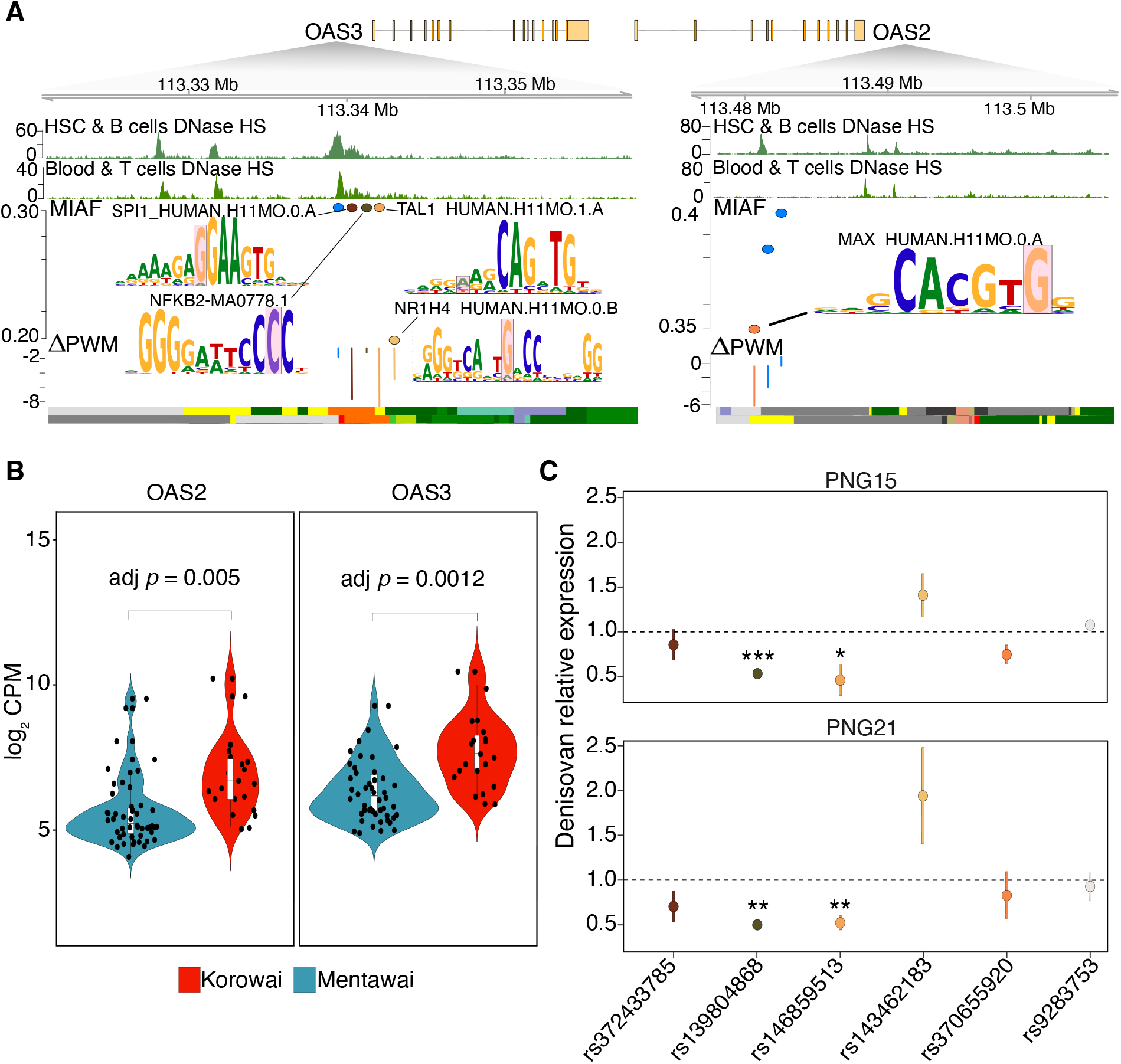
Functional validation of the impact of Denisovan variants on *OAS2* and *OAS3* gene expression. A) Plot of the genomic region encompassing the Denisovan variants associated with *OAS2* and *OAS3*. Top two tracks display patterns of DNase Hypersensitivity sites in Blood and immune T cells as well as in HSC and B cells (Roadmap epigenomics). Tested variants are shown along with their calculated Δ*PW M*. Bottom tracks display the chromatin state for the same tissues. B) log_2_ RNA-seq counts per million (CPM) in whole blood for *OAS2* and *OAS3* between Korowai and Mentawai from [39]. C) Relative expression changes between Denisovan and modern human alleles, or between the alternative and the reference allele for positive control (rs9283753) [43], in two Papuan LCLs. Asterisks mark significant difference from 1.

We tested all five alleles across two different lymphoblastoid cell lines (LCLs) established from Papuan donors. The direction of effect was consistent across biological and technical replicates in all cases. Two Denisovan alleles (rs139804868:A>G and rs146859513:C>G) consistently exhibited significantly lower transcriptional rates than the modern human allele segregating at the same site. rs139804868:A¿G is predicted to disrupt a motif bound by *BHLHA15* and *TAL1*, which have been respectively found to be involved in immune B cell differentiation [40] and hematopoiesis [41]. rs146859513:C>G is predicted to disrupt a binding site for the transcription factor *NFKB2*, which is well known to regulate the expression of a large number of genes involved in both innate and adaptive immune responses, including cytokines and chemokines [42] (Fig. 5c). These observations confirm the validity of our approach and, taken together, our findings systematically point to a substantial contribution of Denisovan variants to immune-related processes in present-day Papuans, one chiefly mediated through the regulation of active immune responses mounted against pathogenic infections.

## Discussion

There is significant interest in understanding the functional consequences of archaic introgression. Evidence indicates that both Denisovan and Neanderthal aSNPs, especially those within protein coding and conserved non-coding elements, are mostly deleterious and negatively selected in modern humans [14, 27]. Similar findings have been recently reported for highly pleiotropic enhancers, where aSNPs are depleted likely as a consequence of their potential to perturb gene expression across multiple tissues [12]. Nevertheless, out of the substantial number of archaic variants still segregating within present-day populations, a large fraction falls within genomic regions that show strong evidence of functional activity across a variety of cell types. Indeed, studies conducted primarily on Neanderthal introgressed DNA have suggested a non-negligible contribution to gene expression variation in modern humans [10, 11, 20], with repeated examples of Neanderthal archaic variants falling within regulatory elements or the seed region of mature micro-RNAs predicted to affect transcriptional and post-transcriptional regulatory processes [11, 44].

In this study, we have taken advantage of a recently published dataset [6] to investigate the landscape of archaic introgression in individuals of Papuan genetic ancestry, the functional consequences of which remained poorly understood. This has allowed us to characterise the putative contribution of Denisovan DNA, which is known to account for up to 5% of the genome of present-day Papuans [4, 45], while also comparing it to that of Neanderthal DNA and regional variants. We specifically analysed all of our variants across multiple cell types and functional chromatin states aiming to account for the strong dependency on the cells’ 1) chromatin landscapes [26]; 2) developmental stages [46] and 3) experienced environmental stimuli [47], when assessing the potential activities of introgressed alleles.

First, we confirm previous reports that aSNPs mostly occur within non-coding sequences [9, 27] with a large number falling within highly constitutive and/or functionally inert genomic regions, which might be the result of weaker purifying selection acting on these elements. However, we find the remaining aSNPs to be significantly depleted from other inactive chromatin states but over-represented within states associated with active gene expression. We also find notable differences between the two archaic ancestries. While Neanderthal and Denisovan variants are both enriched within actively transcribed regions, only Denisovan variants are consistently and significantly enriched within CRE-associated states, and in particular within enhancers. Importantly, variants annotated in these CRE-associated states tend to lie within elements active in a restricted number of cell types, suggesting their biological consequences might be highly cell-type specific. Our findings support the current hypothesis of archaic contributions to pre-and post-transcriptional regulatory processes [11, 44], with important differences among tissues between Neanderthal and Denisovan variants.

Second, we have characterised the impact of introgressed variants on transcription factor binding sites, a well-known mechanism with clear potential to impact phenotypic variation [48]. At least 41.7% of testable SNPs (averaged across all three ancestry sources) falling within CRE-associated states are predicted to modify interactions between TFs and their cognate binding sites. We do not detect significant enrichment or depletion of archaic variants within any cluster of TFs, suggesting that the interplay between TFs and archaic ancestry may be too complex to be fully captured by these whole-genome TFBS footprinting approaches. Therefore, while introgression may have facilitated adaptation by expanding modern humans to new environments [17] our results do not point towards a large role for the rewiring of transcription factor regulatory networks genome-wide as a mechanism. This does not rule out significant impacts on specific TFBSs by any given transcription factor. Indeed, we show that the impact of these TFBS-disrupting variants is not consistent across tissues between the two archaic ancestries.

Third, we consistently find evidence for a sizeable contribution of Denisovan archaic DNA to Papuan regulatory variation, especially within immune-related traits. Only Denisovan high-frequency TFBS-disrupting aSNPs are largely enriched within CRE-associated states active in thymus, HSC & B cells and blood & immune T cells, whereas Neanderthal variants are not enriched in any specific tissue. Again emphasising that different archaic hominins have made different contributions to present-day Papuans, Denisovan and Neanderthal variants predicted to impact gene regulation in immune-related cells generally target different sets of genes, and there is limited overlap in the biological processes they are involved in. Indeed, only genes predicted to be regulated by Denisovan aSNPs are strongly involved in active immune responses, a finding that echoes similar observations made about the role of Neanderthal variants segregating in Europeans [19, 20, 49].

*OAS2* and *OAS3* are among the genes whose regulation we predict likely to be impacted by Denisovan TFBS-disrupting aSNPs. These genes belong to a family of pattern-recognition receptors involved in innate immune responses against both RNA and DNA viruses, with *OAS3* considered to be essential in reducing viral titer during Chikungunya, Sindbis, influenza or vaccinia viral infections [50]. Furthermore, at least two previous studies have shown the presence of both Neanderthal and Denisovan archaic haplotypes segregating at this locus respectively within European [37] and Papuan [38] individuals. In addition, Sams *et al*. found two variants (rs10774671, rs1557866) within these Neanderthal haplotypes which are respectively associated with the codification of different *OAS1* splicing isoforms and with a reduction in *OAS3* expression levels, the latter only upon viral infection [37].

Our previous work has found that both *OAS2* and *OAS3* are differentially expressed between western Indonesians and Papuans [39]. Here we report a set of eight SNPs (rs368816473, rs372433785, rs139804868, rs146859513, rs143462183, rs370655920, rs375463218 and rs372139279), located roughly 41 kb upstream of *OAS2* and *OAS3*, all of which are predicted to strongly alter the ability of different transcription factors, including *IRF4, NFKB2* and *TAL1*, to bind to their underlying DNA motifs. In all cases, the reference allele is fixed within western Indonesian populations, whereas the archaic alleles segregate at frequencies between ≥ 0.2 and 0.4 in Papuans, high enough to suggest an adaptive benefit. We show that at least 5 of these variants lie within sequences that can regulate expression in reporter gene plasmid assays, and that in two of these variants (rs139804868 and rs146859513) the Denisovan allele is associated with significantly lower transcriptional activity compared to the modern human variant in immune cells from two different Papuan donors, again suggesting that these SNPs are of biological importance.

Recent work by [51] has shown that variants which arose in the human lineage following its split from the Neanderthal-Denisovan common ancestor, and which have reached (near) fixation in modern humans since then can exhibit significant differences in regulatory activity relative to the ancestral state. Our results suggest that Denisovan alleles segregating within modern human populations are also likely to actively participate in gene regulatory processes, especially those that take place within immune-related cells. Given the high pathogenic load that characterises the coastal areas of New Guinea island, our findings argue that admixture with archaic hominins might have contributed to immune-related phenotypes amongst early modern humans in the region, favouring adaptation to the local environment [20, 49]. While further high-throughput assays are needed to characterise the genome-wide impact of archaic introgression, our work indicates that admixture with archaic hominins other than Neanderthals might have been a fundamental event in the evolutionary history also for non-European populations. Further decoding of these contributions is essential if we are to improve our understanding of how past events contributed to the phenotypic differences observable among humans today.

## Acknowledgements

We would like to thank all members of the Gallego Romero and McCarthy groups as well as Ashlee Hutchinson (University of Melbourne) for their helpful discussions and comments on the manuscript. We also wish to acknowledge all of the study participants who generously consented to genome sequencing in the original study, and the leadership of Herawati Sudoyo and Chelzie Crenna Darusalam (Eijkman Institute for Molecular Biology) in generating these datasets. This work was supported by an award from the Leakey Foundation and by Australian Research Council Discovery Project DP200101552, both to I.G.R. and by The French National Research Agency (ANR) (grant PAPUAEVOL n° ANR-20-CE12-0003-01 (F.X.R).

## Author contributions

D.M.V. and I.G.R. designed the study. D.M.V. performed the analyses. G.S.J., N.B. and M.P.C. provided raw data. M.L., C.K., F.X.R., N.B. and I.G.R. collected the cell lines used for experimental validation. D.M.V. and L.E.C. designed the plasmid reporter assay experiment. D.M.V. performed the experiment and analysed the data. L.E.C. provided feedback on analysis of the plasmid reporter assay data. D.M.V. and I.G.R. wrote the paper with input from all other authors. All authors approved the manuscript before submission.

## Materials and Methods

### Samples

All human genome data in this study was taken from Jacobs *et al*. [6]. All collections followed protocols for the protection of human subjects established by institutional review boards at the Eijkman Institute (EIREC #90) and Nanyang Technological University (IRB-2014-12-011); the analyses in this publication were additionally approved by University of Melbourne’s Human Ethics Advisory Group (1851585.1). Permission to conduct research in Indonesia was granted by the State Ministry of Research and Technology (RISTEK). All individuals gave their full informed consent to participate in the study.

The two lymphoblastoid cell lines used in the plasmid reporter experiments were established from donors sampled at the University of Papua New Guinea, in Port Moresby, Papua New Guinea. Collection of material from healthy adult donors was coordinated by Dr Christopher Kinipi (Director of Health Services at the University of Papua New Guinea) and approved by the Medical Research Advisory Committee of the National Department of Health of the Government of Papua New Guinea (permit number MRAC 16.21), by the University of Melbourne (ID: 1851585.1), and by the French Ethics Committees (Committees of Protection of Persons 25/21_3, n°SI:21.01.21.42754). Permission to conduct research in Papua New Guinea was granted by the National Research Institute of Papua New Guinea (permit 99902292358), with full support from the School of Humanities and Social Sciences, University of Papua New Guinea. These approvals commit our team to following all ethical guidelines mandated by the government of Papua New Guinea.

### Variant filtering and selection

To curate a high confidence set of SNPs segregating in our dataset, we began by intersecting genome-wide genotype data from all 56 genetically Papuan individuals with the set of high confidence Neanderthal and Denisovan haplotypes identified in this group, as defined by [6] (for detailed methods and sample information see [6]). Due to incomplete lineage sorting, these regions may still contain SNPs that are not Denisovan or Neanderthal-specific, thus we applied a series of stringent filtering steps, as follows. Variants were initially labelled as aSNPs or naSNPs depending on whether they fell within archaic haplotypes, and all singletons, regardless of background, removed. Only SNPs lying outside both sets of archaic haplotypes were considered potential naSNPs, with the remaining variants were sorted into Denisovan aSNPs or Neanderthal aSNPs based on the inferred ancestry of the haplotype of origin. aSNPs in common between the two archaic haplotypes were assigned, where possible, to one of the two ancestries by considering whether the state of the main introgressed allele in Papuans resembled the homozygous state of the Altai Neanderthal [52] or of the Altai Denisovan [53] reference genomes. Variants that could not be disambiguated in this manner were labelled as ambiguous and excluded from downstream analyses. We further excluded from our analyses all aSNPs where both alleles were present at identical frequencies within archaic haplotypes, impeding identification of the main introgressed allele, as well as naSNPs for which the ancestral allele is fixed within non-archaic haplotypes. Finally, we removed all SNPs, both of putative archaic and modern ancestries without ancestral state information in the 6 different non-human primates EPO alignment.

Allele frequencies were calculated by considering the fraction of Papuan individuals carrying the variant within the haplotype of the respective ancestry. For naSNPs, we calculated the derived allele frequency (DAF) by dividing the number of observations of the derived allele in individuals carrying the modern haplotype over the total number of chromosomes at that locus. Likewise, for each aSNP, we defined the major introgressed allele frequency (MIAF) as the number of observations of the major allele in individuals carrying an archaic haplotype, divided by the total number of chromosomes. For each aSNP we also calculated the MIAF in individuals carrying a modern human haplotype, which we later used to filter our variants (see below). Variants with MIAF/DAF < 0.05 were labelled as low-frequency and removed from downstream analyses.

We then applied two additional filters to create a high-confidence set of aSNPs. To control for incomplete lineage sorting between humans, Neanderthals and Denisovans [2], we first computed the absolute difference between the frequency of the main introgressed allele between archaic and modern human haplotypes and discarded all aSNPs where this difference was ≤ 0.25. Second, we removed all aSNPs for which the main introgressed allele is segregating at frequencies ≥ 0.005 within sub-Saharan African populations from the 1000 Genomes Project [54]. We also applied this filtering criteria to the non-archaic derived alleles to ensure a fairer comparison between naSNPs and aSNPs. Following these steps, we retained 79,808 Denisovan, 51,268 Neanderthal and 338,585 non-archaic alleles, which were used in all downstream analyses.

### SNP annotation and enrichment across chromatin states and cell types

We downloaded the 15-state hg19 mnemonics BED files containing cell type-specific segmented chromatin state predictions for 111 cell types from the Roadmap Epigenomics project [26], https://egg2.wustl.edu/roadmap/web_portal/index.html. SNPs were intersected with this chromatin state information to define their potential activity across every cell type. Likewise, we downloaded the list of V8 expression quantitative trait loci (eQTLs) and their eGenes from the GTEx web portal https://www.gtexportal.org/home/ [28]. Genomic coordinates were lifted over from hg38 to hg19 using liftOver from the rtracklayer R package [55] and only significant eSNP-eGene pairs (FDR *q*-value ≤ 0.05) were retained. We then merged the two lists of variants, retaining only those instances where both the reference and alternative alleles matched between the two datasets.

Throughout the study, to quantify aSNPs’ impact within each chromatin state and/or across each chromatin state/cell type combination, for each ancestry, we counted the number of variants annotated in each combination of interest. We then computed the odds ratio (OR) between the number of aSNPs and naSNPs for Denisovan and Neanderthal separately and tested eventual significant deviations from an OR = 1 via Fisher’s exact test (all statistical results can be found in Supp Table 1. Reported values have been corrected for multiple testing as in [56].

To assess whether SNPs fell within highly pleiotropic, constitutive or cell-specific functional elements, we grouped the 111 cell types into the 18 tissues as reported by [26], and for each ancestry we counted across how many of these SNPs are annotated to a particular chromatin state. For each chromatin state, we then calculated the cumulative proportion of SNPs having a particular pleiotropic range [1, 111].

### Variant impact on TFBSs

To characterise the impact of our sets of SNPs on transcription factor binding sites (TFBSs), we focused on variants that fell within TssA, TssAFlnk, TxFlnk, Enh and EnhG in at least one cell type, and defined the union of these regions as putative cis-regulatory elements (CREs). To assess whether any of our SNPs might disrupt any TFBS we used the motifbreakR R package (v 2.2.0) [57], retrieving the collection of DNA motifs from Jaspar 2018 [30] and HOCOMOCO v.11 [29] databases. Motifs were matched to the hg19 genome using a *p*-value threshold of 1^-5^, and the allelic impact on each identified motif was quantified via the built-in sum of log-probabilities algorithm, setting a background probability of 0.3 and 0.2 for A/T and G/C, respectively. Allelic impact on the position weight matrices (PWMs), i.e. Δ PWM score, was obtained from motifbreakR. However, because motifbreakR reports ΔPWM as the difference in PWM scores between the reference and alternative allele, we re-polarised such score with respect to the introgressed relative to the non-introgressed allele (for aSNPs) or with respect to the derived relative to the ancestral allele (for naSNPs). Duplicated variants predicted to disrupt the same binding site across both databases were removed by retaining only the SNP-PWM association with the highest absolute Δ PWM score. The final set of SNPs, together with their affected motifs and their predicted disruptive potential, are listed in Supp File 1 -Supp File 3.

To assess the genome-wide impact of aSNPs on TFBSs, and to avoid redundancy between TFs due to similar binding preferences across motifs, we grouped all our TFs into families based on motif clustering information from [31]. We removed 72 HOCOMOCO and 93 Jaspar PWMs that were either not associated with any cluster or that have mismatches in the PWM versions between MotifDb [58] and [31]. For each cluster we then computed the OR between the associated number of Denisovan or Neanderthal aSNPs relative to that of naSNPs, testing any eventual deviations from an OR = 1 via Fisher’s exact test and applying FDR-correction to resulting *p*-values (Supp Table 2).

Finally, to evaluate the impact of both archaic and modern human derived alleles on their TFBSs we tested whether the distribution of ΔPWM scores for these variants was significantly different from 0 across each of the 18 tissue types. Resulting *p*-values were then FDR-corrected since TFBS-disrupting SNPs can be annotated over multiple tissues.

### GO enrichment analysis

To understand the biological processes potentially affected by Denisovan and Neanderthal introgression, we retrieved all the TFBS-disrupting variants annotated within CREs active in at least one HSC & B and/or blood & T related cell type. We then used the rGREAT R package (v 1.20.0) [59] to assign these variants to the regulatory domain of the two nearest genes located within 1 megabase of distance in either direction from the genes’ transcription start sites. For each ancestry, we independently performed a GO enrichment analysis [60], using as a background set the union of all the TFBS-disrupting variants associated genes (both archaic and non-archaic) (Supp Table 3).

To quantify similarity between enriched GO terms across all ancestries, we computed pairwise semantic similarity scores across all three ancestry pairs using the Wang similarity metric as implemented in the GOSemSim R package [61]. Briefly, this metric quantifies the semantic similarity of each pair of GO terms based on both their locations in the GO graph as well as their relationship with their ancestor terms (see [35]). The resulting matrix was then summarised into a single value using the Best-Match-Average (BMA) method, which calculates the average of all maximum similarity scores across GO term ID comparison.

### Plasmid reporter assays

We designed 12 170 bp oligonucleotides centred on the archaic and modern alleles of the 5 *OAS* SNPs discussed above (84 bp on the 5’ and 85 bp on the 3’ end), as well as for rs9283753, a control site from Tewhey *et al*. 2016 [43]. All variants were on the positive strand, regardless of variant orientation with respect to its predicted target gene, as enhancer activity in reporter assays has been found to be largely independent of orientation, [62]. In addition, all oligos contained two common 15 bp adapter sequences flanking each regulatory element, a 44 bp sequence centred on a BsiWI restriction site, a oligo-specific 15 bp barcode sequence and a universal 20 bp buffer sequence, resulting in a final oligo length of 284 bp. Oligos were synthesised as a pool by IDT. Full sequences of all oligos are available as Supp Table 4.

Synthesised oligos underwent two consecutive runs of Q5 (NEB, M0491) low-cycle PCR using two different sets of primers (see Supp Table 4), first to make them double-stranded and then to add a SfiI restriction site at each end of the sequence. Following each PCR amplification, oligos were purified with AMPure XP beads (Beckman Coulter, A63881). Amplified oligos and pMPRA1 reporter plasmid (Addgene, 49349) were both SfiI digested (NEB, R0123), ran on a 1.5% agarose gel, extracted (NEB, T1020) and ligated using T4 DNA Ligase (NEB, M0202) always following manufacturer’s protocols. 50 ng of ligated product was transformed into 25 μl of *E.coli* competent cells (Promega, JM109), recovered in 250 μl of SOC for 1 hour at 37 °C following heat shock. Bacterial cultures were then grown overnight in 25 mL of LB media supplemented with 100 μg/mL of ampicillin (Thermo Fisher Scientific, FSBBP1760-5) on a floor shaker at 37 °C, prior to plasmid purification (Zymo, D4209).

To create the final library, assembled plasmids were linearised with BsiWI-HF (NEB, R3553). An amplicon containing a minimal promoter, GFP open reading frame (ORF), previously synthesised by IDT (Supp Table 4), was then inserted by Hi-Fi assembly (NEB, E2621) using 0.025 pmols of vector and 0.05 pmols of the GFP amplicon, incubating at 50 °C for 1 hour. 50 ng of assembled product were then transformed into bacterial cells, which were then grown overnight at 37 °C and subsequently purified (Zymo, D4200). For each step, the transformation efficiency was estimated to be > 10^6^ cfu. To ensure correct assembly, approximately 150 ng of purified products were first ran on a 1.5% agarose gel, extracted, PCR-amplified using an universal forward primer binding to the 5’ adapter sequence and an oligo-specific reverse primer binding to the 3’ barcode sequence, and confirmed by Sanger sequencing (Macrogen).

LCLs for experimental validation were grown in RPMI (Lonza, 12-702Q) supplemented with 1% Glutamax, 1% NEAA and 10% FBS. Cells were maintained at a cell density of 5 · 10^5^ cells/mL. Each cell line was electroporated in triplicate using the Neon transfection system (Thermo Fisher Scientific, MPK5000). For each transfection, 1 · 10^6^ cells were centrifuged at 700x g and resuspended in 100 μl of R buffer with 1 μg of plasmid library before applying 3 pulses of 1200 V for 20 ms each [43]. Following transfection, cells were allowed to recover in 2 mL of culture media for 24 hours then collected by centrifugation and frozen at −80°C.

For each transfection replicate, total DNA, including plasmid DNA (pDNA), and RNA were extracted from cells using Qiagen AllPrep kit (Qiagen, 80204) following the manufacturer’s protocol, including on-column DNase digestion of the RNA fraction. pDNA and total RNA samples were quantified using the Qbit broad range dsDNA (Thermo Fisher Scientific, Q32850) and RNA kits (Thermo Fisher Scientific, Q10211). First-strand cDNA was synthesised from 1 μg of mRNA for each sample using the High-Capacity cDNA Reverse Transcription Kit (Applied Biosystems, 4368814) with custom primers (Supp Table 4).

The regulatory activity of each oligo was then quantified via qrtPCR. We first assessed relative primer binding efficiencies through a series of 4 (1:10) serial dilutions of an aliquot of cDNA. These efficiencies were incorporated in the calculation of each oligo’s relative expression (see below). For each transfection replicate we quantified the relative abundance of pDNA and cDNA for each oligo using a common forward primer binding within the GFP ORF and 12 different reverse primers, each targeting the oligo-specific barcode sequence. All reactions were performed in triplicate. Relative expression of each oligo for each transfection replicate was calculated with the Δ*C_t_* method, normalising it by the relative amount of pDNA. Expression differences between archaic and modern human alleles were calculated using the relative expression ratio [63].

## Code availability

All analyses were performed using R software v 4.0.0. The complete list of scripts used for the analyses is available at https://github.com/dvespasiani/Archaic_introgression_in_PNG

## Supplementary materials

Supplementary materials include 5 figures, 4 tables and 3 files.

**Supp Table 1** Table reporting the aSNPs OR results across each 1) chromatin state; 2) chromatin state/cell type combination; for the set of 3) all and 4) New Guinean-specific TFBS-disrupting variants

**Supp Table 2** Table reporting 1) the aSNPs OR across TF clusters; 2) the *p*-values associated with the SNPs impact on PWMs across tissues.

**Supp Table 3** Table reporting the GO enrichment results for both aSNPs and naSNPs associated genes

**Supp Table 4** Table reporting the primer and oligo sequences used in the ***in-vitro*** validation.

**Supp File 1** File reporting all Denisovan TFBS-disrupting aSNPs

**Supp File 2** File reporting all Neanderthal TFBS-disrupting aSNPs

**Supp File 3** File reporting all modern TFBS-disrupting naSNPs

## Supplementary Materials

**Fig S1.**
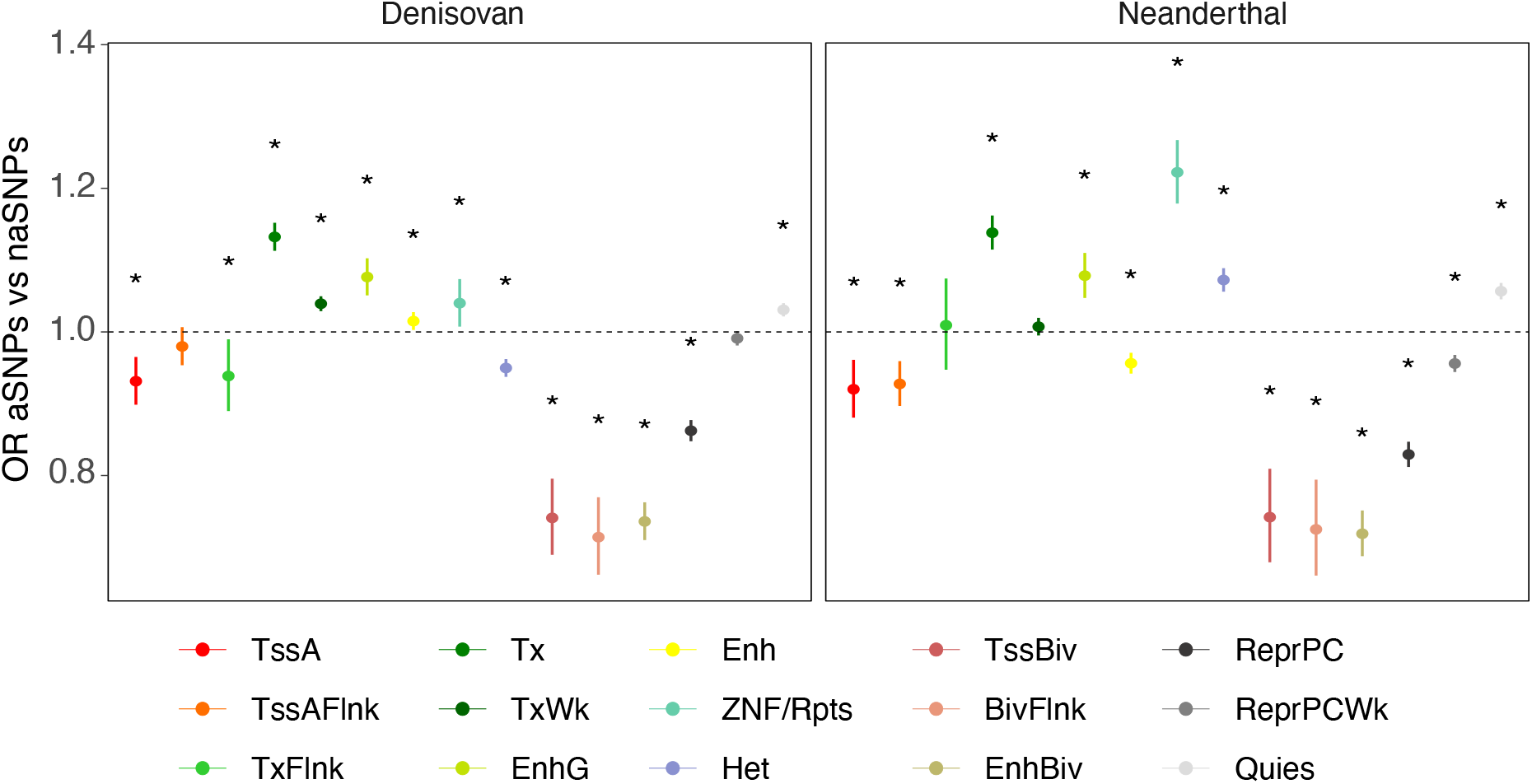
aSNPs enrichment across 15 chromatin states. Denisovan and Neanderthal enrichment for each chromatin state. Dots and bars represent OR and upper/lower range of confidence interval. Asterisks indicate FDR-corrected Fisher’s exact test *p* < 0.05 (see Supp Table 1)

**Fig S2.**
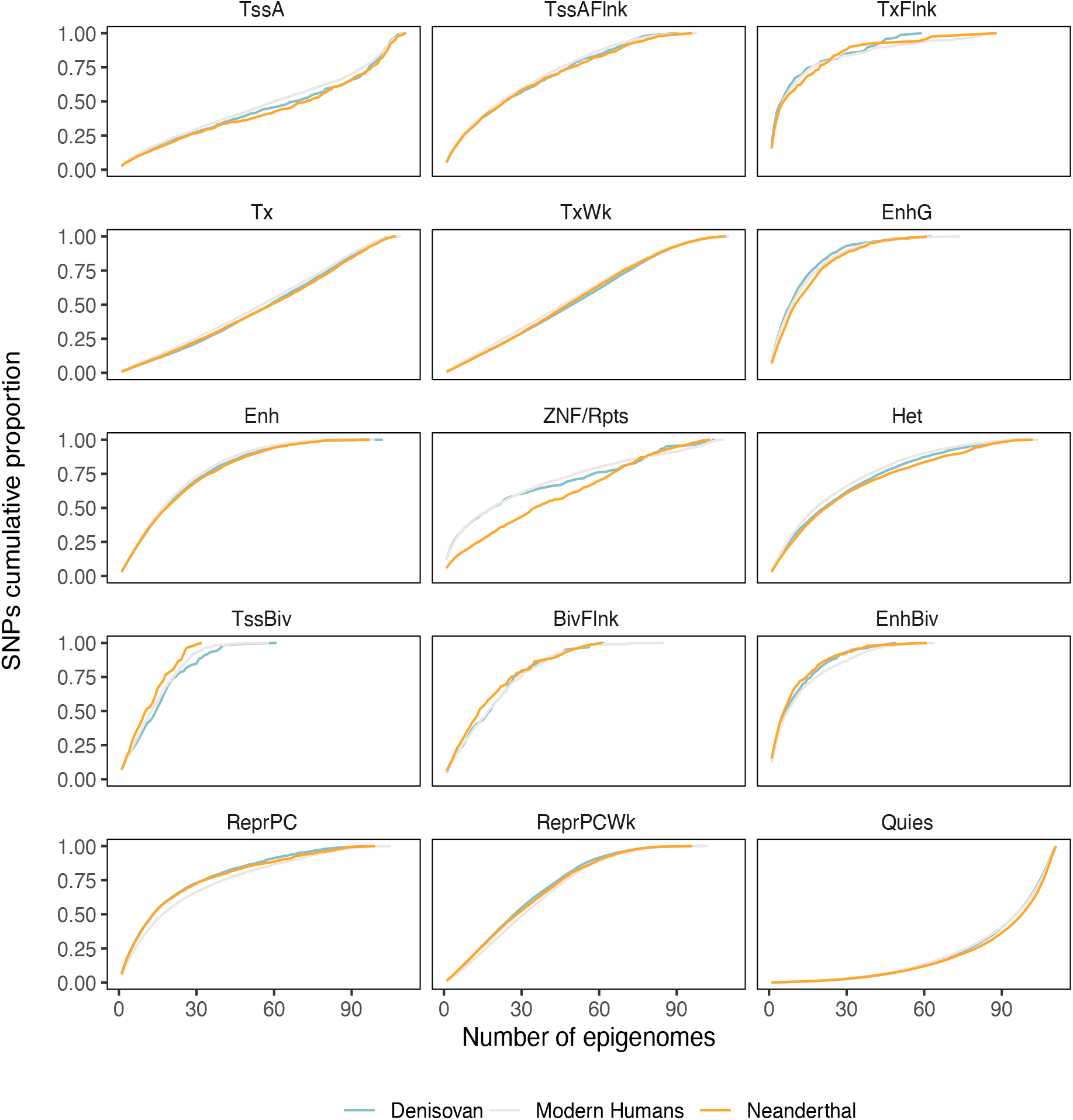
Pleiotropic activities of chromatin state-associated elements carrying SNPs. Cumulative proportion of the pleiotropic activity across 111 cell types of each chromatin state-associated element carrying SNPs.

**Fig S3.**
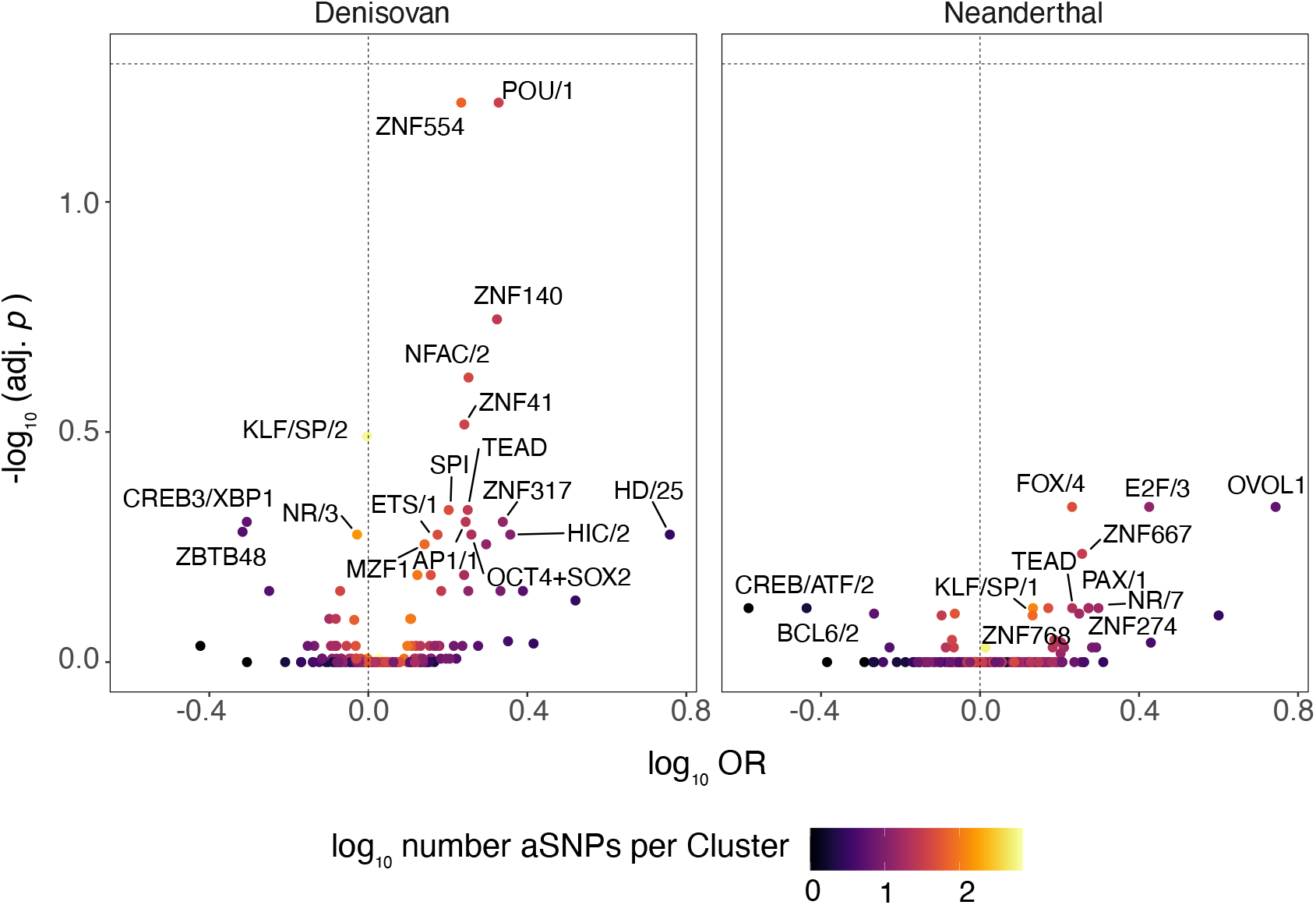
aSNPs enrichment across TF clusters. Enrichment of Denisovan and Neanderthal TFBS-disrupting variants across TF clusters. Dots are coloured based on the number of archaic variants annotated within the given cluster. Only clusters with a nominal *p* < 0.05 are labelled.

**Fig S4.**
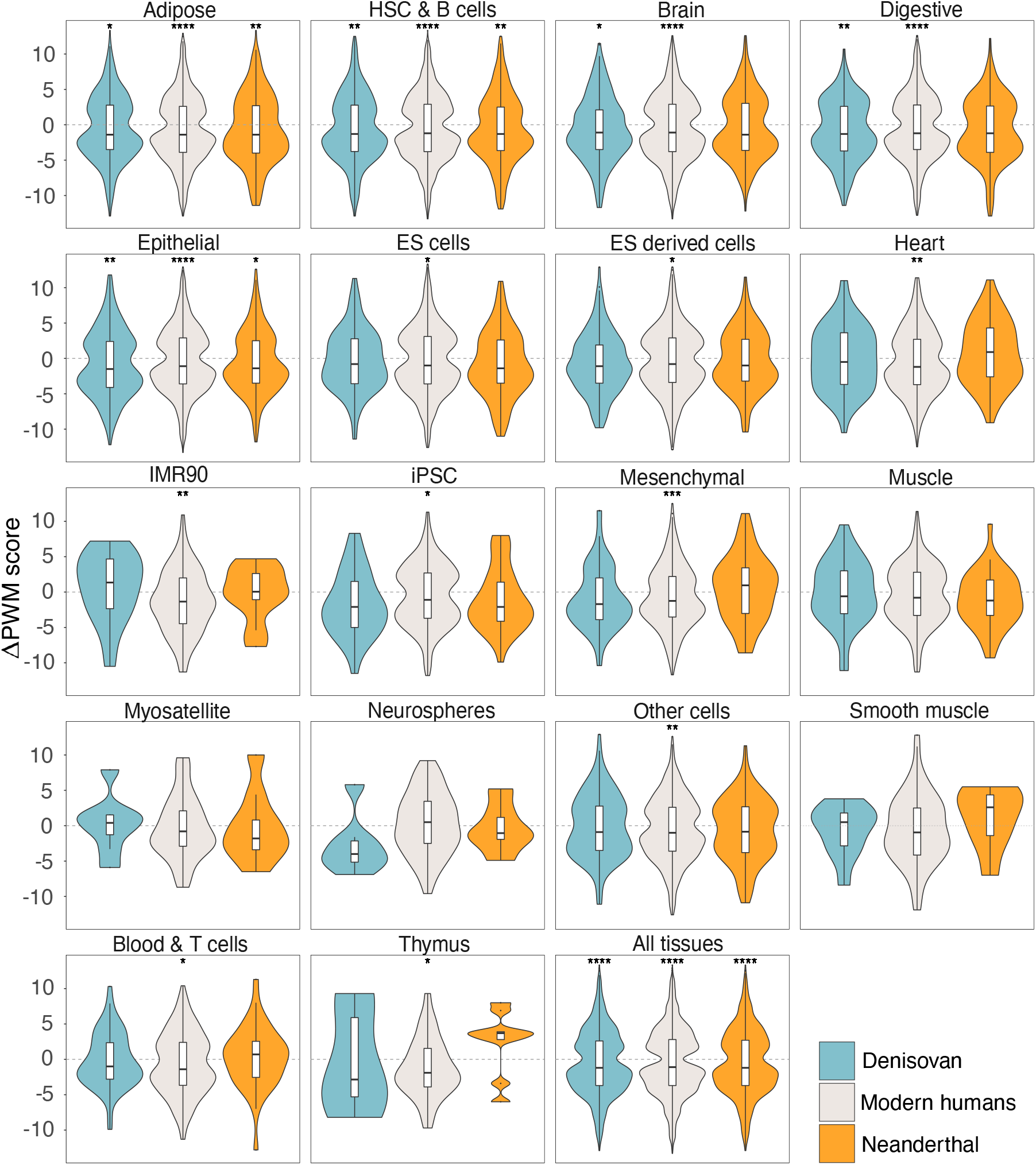
Impact of TFBS-disrupting alleles on DNA motifs across tissues. The calculated ΔPWM score with respect to the main introgressed/modern human derived allele, respectively for the set of high-frequency TFBS-disrupting aSNPs and naSNPs. Dashed line corresponds to Δ*PW M* = 0. Violin plots represent the full distribution of the ΔPWM scores for the sorted cell types. Lower and upper hinges of the boxplots correspond to the first and third quartiles of the distribution, whiskers extend to a maximum of 1.5 × IQR beyond the box. Asterisks indicate whether the median ΔPWM scores is significantly different from 0 (FDR-adjusted one-tailed Wilcoxon rank-sum test, see Supp Table 2).

**Fig S5.**
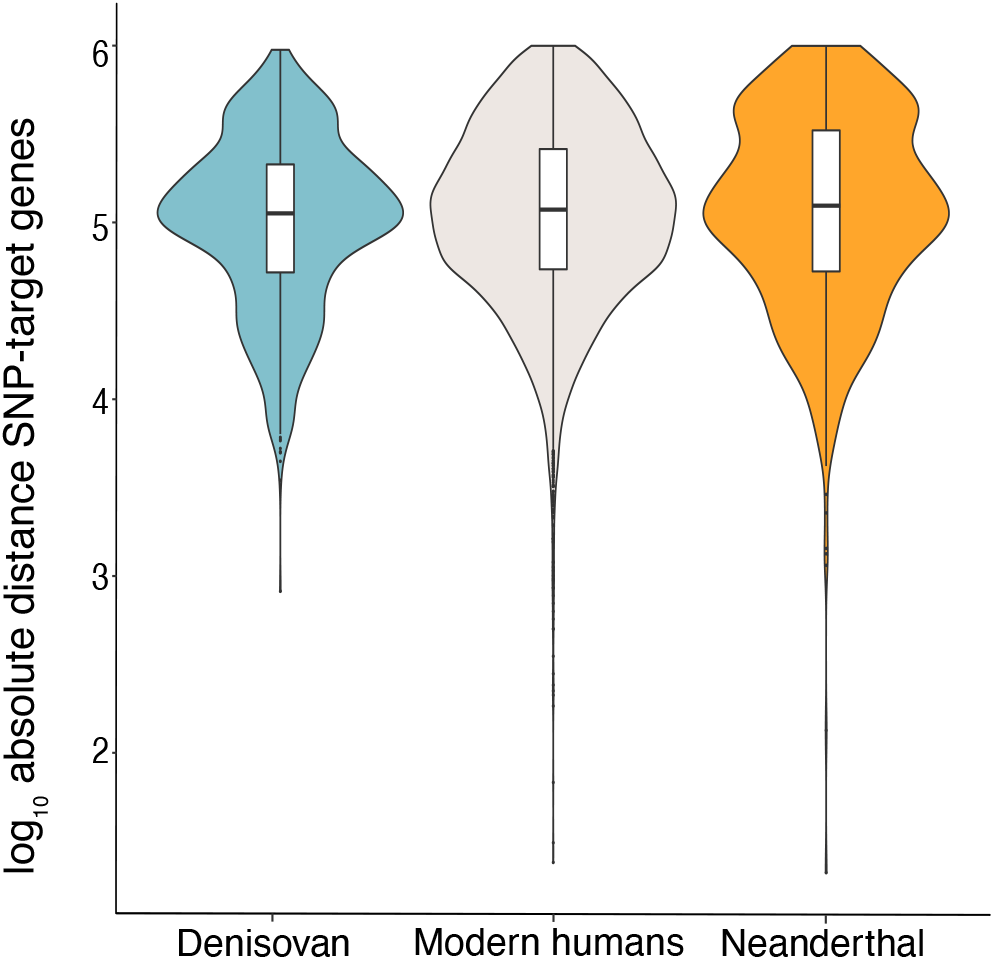
SNPs-targe gene distance. The distance in log_10_ bp between the SNPs and their target genes as predicted by GREAT [34]. Violin plots represent the full distribution of the log_10_ bp distances. Lower and upper hinges of the boxplots correspond to the first and third quartiles of the distribution, whiskers extend to a maximum of 1.5 × IQR beyond the box.

